# Dissociation of the mTOR protein interaction network following neuronal activation is altered by Shank3 mutation

**DOI:** 10.1101/2025.05.20.655155

**Authors:** Devin T. Wehle, Emily A. Brown, Vera Stamenkovic, Felicia Harsh, Stephen E.P. Smith

**Author notes:** Correspondence to: S.E.P.S.

## Abstract

The mechanistic target of Rapamycin (mTOR) kinase pathway plays critical roles in neuronal function and synaptic plasticity, and its dysfunction is implicated in numerous neurological and psychiatric disorders. Traditional linear models depict mTOR signaling as a sequential phosphorylation cascade, but accumulating evidence supports a model that includes signaling through dynamic protein-protein interaction networks. To examine how neuronal mTOR signaling discriminates between distinct stimuli, we quantified phosphorylation events and protein co-association networks in primary mouse cortical neurons. Unexpectedly, neuronal mTOR activation by IGF or glutamate triggered dissociation—rather than the anticipated assembly—of protein complexes involving mTOR complex1 (TORC1), mTOR complex 2 (TORC2), and translational machinery, distinguishing neurons from proliferative cells. Applying in vitro homeostatic scaling paradigms revealed distinct combinatorial encoding of synaptic scaling direction: both up- and down-scaling induced dissociation of translational complexes, but downscaling uniquely included dissociation of upstream pathway regulators. Cortical neurons from Shank3B knockout mice, modeling autism-associated Phelan-McDermid Syndrome, displayed baseline hyperactivation of the mTOR network, which reduced the dynamic range of network responses to homeostatic scaling and pharmacological inhibition. These findings reveal that neuronal mTOR signaling employs stimulus-specific combinations of dissociative protein interaction modules to encode opposing forms of synaptic plasticity.

## Introduction

Eukaryotic cells rely on intricate signal transduction networks to process and respond to their external environment. Traditional linear cascade models, exemplified by kinase signaling pathways such as the MAPK cascade (*1*), describe signals transmitted through sequential phosphorylation events that relay information in a direct and specific manner. However, evidence demonstrating that signaling proteins often physically interact in highly interconnected networks challenges the simplicity of this linear paradigm. An alternative model, proposed by Tony Pawson following discovery of the SH2 domain, posits that signaling proteins form dynamic complexes, with specific protein combinations encoding stimulus-specific information (*2*). These two signaling models are not mutually exclusive, and each has substantial experimental and theoretical support (*3*, *4*). Reconciling these two models into a cohesive theory remains challenging, since the linear framework implies both direct causality and specificity, but protein interaction networks are highly interconnected, and interactions are often promiscuous.

mTOR is a serine/threonine kinase critical for regulating eukaryotic cell growth, proliferation, and nutrient-responsive metabolism (*5*). In neurons, which do not proliferate and live in a relatively stable nutrient environment, mTOR signaling has been adapted for specialized processes, including neuronal-activity-dependent protein synthesis, calcium homeostasis, autophagy, and synaptic plasticity (*6*). Given the central importance of these diverse processes to neuronal function, it is perhaps not surprising that mTOR dysregulation has been implicated in a vast number of neurological disorders, including epilepsy (*7*), brain overgrowth disorders (*8*) and neuropsychiatric disorders including autism (*9*, *10*). In fact, haploinsufficiency of several components of the mTOR pathway are related to syndromic forms of autism (*11*, *12*), and data from mouse models and human postmortem studies (*10*, *13*, *14*) indicate that mTOR dysregulation is a recurring feature of both genetic and idiopathic autisms.

mTOR signaling is often modeled as linear pathway, in which PI3K activation leads to AKT phosphorylation, which leads to mTOR phosphorylation and then phosphorylation of pathway outputs S6K and 4E-BP1, associated with translation (*15*). However, direct protein-protein interactions among mTOR pathway components suggest a more complex network architecture. For example, upstream PTEN interacts directly with downstream EIF4E (*16*), indicating wiring beyond the linear pathway.

Perhaps a better model of mTOR system-level behavior would help to reconcile ongoing debates about the function of specific mTOR components (*17*), or improve drug normalization of phenotypes caused by hyperactivating mutations in PI3K, PTEN, TSC2 or mTOR (*18*).

To that end, we recently compared linear vs. network models of the mTOR system by simultaneously monitoring phosphorylation events by phospho-western blot, and protein complex composition by quantitative multiplex co-immunoprecipitation (QMI). We applied various small molecule inhibitors during stimulation to immortalized fibroblasts, and found that while phosphorylation events progressed as predicted from PI3K to AKT to mTOR, protein complexes increased their co-associations in a modular, stimulus-specific pattern that did not correspond to a linear hierarchy (*19*).

Here, we extend this analysis to primary mouse cortical neurons, assessing both mTOR phosphorylation and co-association networks to elucidate how mTOR differentiates neuronal signals. We find that mTOR signaling distinguishes opposing types of plasticity-inducing synaptic inputs through a modular and combinatorial encoding mechanism, and that the dynamic range of this signaling system is disrupted in Shank3B knockout mice, a genetic model of Phelan-McDermid syndrome. Moreover, we show that the relationship between mTOR network behavior and phosphorylation status differs substantially between neurons and fibroblasts, which suggests cell type-specific rules for integrating linear and network-based signaling models.

## RESULTS

### mTOR activation associated with network dissociation in neurons

We previously demonstrated that growth factor starvation and refeeding of immortalized mouse fibroblasts caused a relatively slow (∼1-hour), coordinated increase in mTOR network protein-protein interactions (19). To determine if the mTOR network behaved similarly in mouse neurons, we subjected DIV 18-21 mouse cortical neurons to 2 hours of growth factor starvation, followed by stimulation with IGF1 or glutamate, both known to activate mTOR signaling (*20–22*) (Figure 1A). Phospho-AKT was reduced by growth factor starvation (removal of B27 supplement), and increased with exposure to IGF1 or glutamate (Fig 1B,C). We used quantitative multiplex co-immunoprecipitation (QMI, see methods section for details) to simultaneously monitor 306 protein interactions among 18 mTOR pathway proteins, using a QMI antibody panel previously validated in mouse tissue (Table S1). Principal components analysis on the QMI data matrix (consisting of 306 interactions monitored across 4 conditions, N=4 biological replicates per condition) revealed that growth factor starvation acted to compact the biological replicates in PCA space compared to untreated “baseline” neurons, reducing variability due to stochastic fluctuations in mTOR activity (Fig 1D). IGF1 and glutamate stimulation overlapped with each other, but not with baseline or starvation, suggesting a similar network response to both stimulus types.

**Figure 1:**
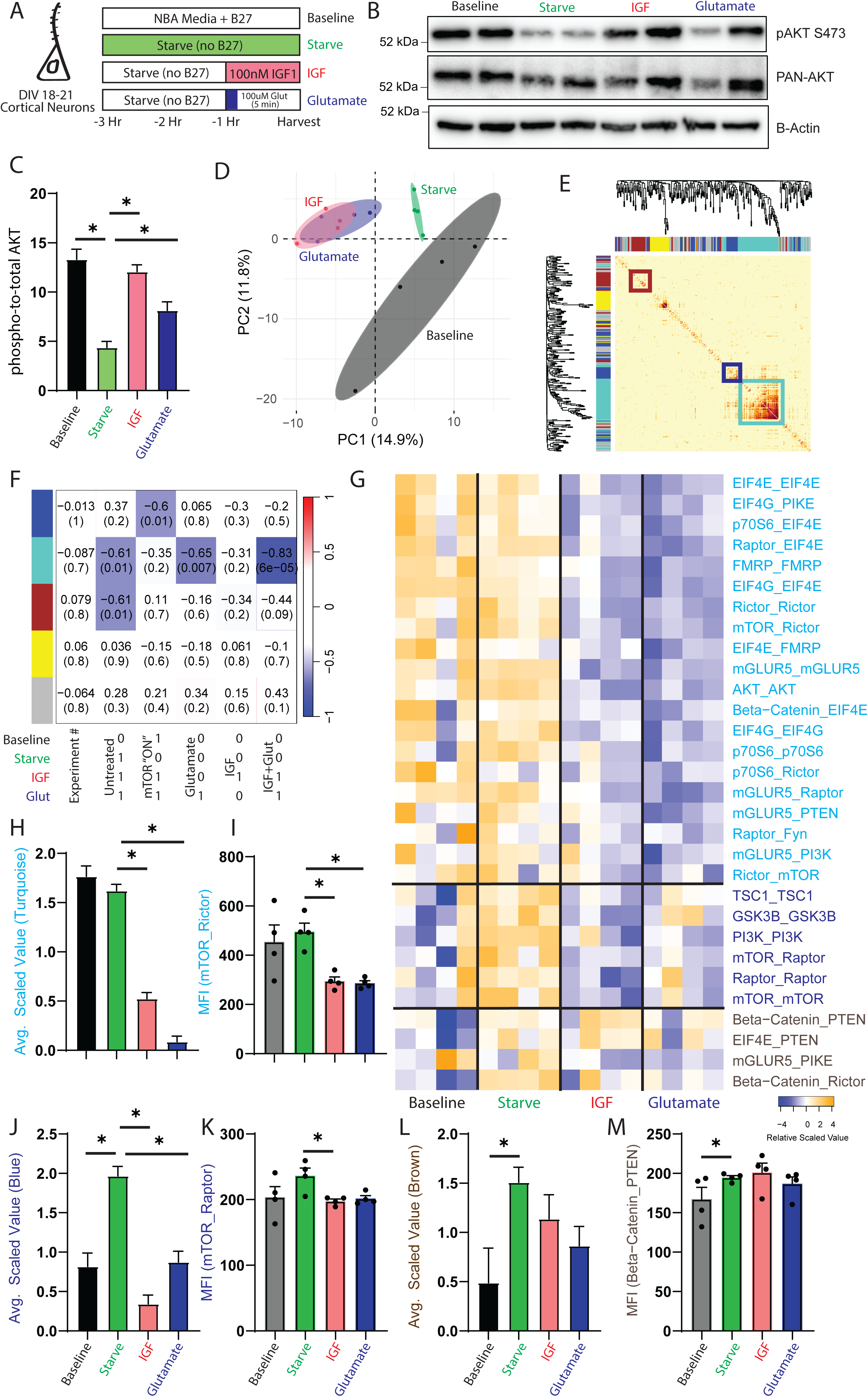
mTOR dynamics following IGF or Glutamate stimulation. A) Experimental design. NBA, neurobasal media; DIV days in vitro. B) Representative western blots showing phospho- and total AKT, with actin for a loading control. C) Quantification of blots shown in B. * indicates p<0.05 from starved by ANOVA followed by Dunnett post-hoc testing,. N = 4. D) Principal component graph of QMI data, N=4 per condition. E) Topological overlap matrix of QMI data. Ends of the symmetrical dendrogram indicate interactions (unlabeled). Dark pixels indicate stronger correlation between interactions on the x and y axes. Colored boxes indicate modules. F) Module-trait table showing the correlation coefficient (top number) and p-value (bottom number) between the eigenvector of each color-coded module (colored rectangles on the left) and binary-coded variables as shown in the table below. G) Heatmap of scaled values of all significantly changed interactions. Each box represents a single interaction measurement from a single biological replicate; columns correspond to a biological replicate while rows correspond to an interaction (listed IP_Probe). Statistical significance calculated by ANC and CNA statistics as detailed in methods. H-M) Mean scaled value of all interactions in the turquoise (H), blue (J) or brown (L) module, and a representative interaction from each module (I,K,M). * indicates p<0.05 by ANOVA followed by Dunnett post-hoc testing (H,J,L) or by ANC (I,K,M).

Protein interaction networks have evolved a modular organization, which increases robustness and decreases noise inherent in individual interactions (*23*, *24*). We used correlation network analysis (CNA) (*25*) to cluster interactions into modules based on similar behavior across the 16 samples, visualized as a Topological Overlap Matrix (TOM) plot (Figure 1E). Three of these modules-arbitrarily named “blue”, “turquoise” and “brown”, correlated with binary-coded experimental variables representing stimulation states (Fig 1F). The largest module, “turquoise”, best correlated with a model in which its constituent interactions dissociated following IGF1 or Glutamate stimulation (correlation coefficient = -0.83, p = 6 x 10^-5^), suggesting “turquoise” encodes mTOR activation. The “blue” module correlated with a model in which interactions dissociated in all conditions except for serum-starved (correlation coefficient = -0.6, p = 0.01), suggesting this module reflects tonic mTOR activation that was relieved by starvation and reinstated by IGF or Glutamate. Finally, the “brown” module best correlated with a model in which co-associations decreased following starvation and did not return to baseline after either treatment (correlation coefficient = -0.61, p = 0.01).

To identify the pairwise interactions that comprise each module, we used an <u>A</u>daptive, <u>N</u>on-parametric, paired statistical test <u>C</u>orrected for multiple comparisons (ANC), which was specifically designed for QMI datasets (*26*). Interactions that were both significant by ANC (Bonferroni-corrected p<0.05) and a member of the turquoise, blue or brown CNA modules were plotted on a row-normalized heatmap in descending order of module correlation strength (Fig 1G). The IGF-and-Glutamate-responsive “turquoise” module consisted of 20 interactions whose averaged scaled value was not altered by starvation, but was significantly *reduced* by both IGF and glutamate stimulation (Fig 1H). The most prominent interactions involved eukaryotic initiation factor (EIF)4E and EIF4G, which are components of the translation initiation complex downstream of the mTOR pathway. IGF and glutamate treatment each strongly reduced the median fluorescent intensity (MFI) of multiple interactions involving EIG4G and EIF4E in shared complexes with RAPTOR, P70S6 kinase, and FMRP. Additionally, IP:mTOR probe:Rictor (abbreviated mTOR_Rictor, Fig 1I), Rictor_mTOR and Rictor_Rictor were reduced by stimulation, indicating changes to mTOR complex 2 (TORC2). We validated the reduction in IP:mTOR probe Rictor by IP-western blot and found a similar reduction in the amount of co-associated protein (Fig S1).

The starvation-activated “blue” module consisted of 6 interactions that increased in abundance during starvation and returned to the basal (dissociated) state upon stimulation (Fig 1J). Prominent among the blue module were the mTOR complex 1 (mTORC1) interactions mTOR_RAPTOR (Fig 1K), RAPTOR_RAPTOR and mTOR_mTOR, as well as PI3K, GSK3b and TSC1 self-detection (indicating either decreased abundance, self-association, or epitope availability). Finally, four brown-module interactions increased with starvation but did not return to baseline by 1 hour (Fig 1L), including Beta-Catenin_PTEN (Fig 1M), and mGluR5_PIKE. Overall, these data indicate that the response to mTOR-activating stimuli in neurons is the dissociation of protein interactions representing all levels of the mTOR signaling cascade: TORC1, TORC2 and translational outputs. This was surprising, because in fibroblasts mTOR activation and phosphorylation correlated with increased, not decreased, co-associations (*19*).

### mTOR activation in Shank3 neurons

The autism risk gene Shank3 contributes to the integrity of glutamatergic post-synaptic receptor signaling complexes, and the loss of Shank3 disrupts glutamatergic transmission and reduces activation of downstream signaling pathways such as Erk and mTOR (*27*, *28*). To characterize mTOR network behavior in Shank3 deficient neurons, we starved DIV 18-21 cultured cortical neurons from Shank3B knockout mice (*28*), or their wildtype littermates, and stimulated them with IGF1 or Glutamate as above. We did not use the ‘baseline’ condition because it was similar to, but more variable than, the starved condition (Fig 1D). Western blots showed phosphorylation of AKT following IGF1 treatment, while Glutamate treatment yielded a small non-significant response in both genotypes (Fig 2A,B). QMI analysis identified a “green” module that correlated with IGF and Glutamate stimulation (correlation coefficient = -0.36, p = 0.03). Importantly, 14/17 interactions in the green/stimulation module were independently identified in the experiment shown in Figure 1 (bolded in Fig 2C), demonstrating robust detection of responsive protein complexes. A heatmap of significant interactions suggested that the Shank3B^-/-^ neurons did not respond as strongly as WT neurons (Fig 2C). Indeed, while the averaged scaled value of the module was significantly reduced in WT neurons in response to both IGF and Glutamate, there was no significant reduction in Shank3B^-/-^ neurons in response to IGF treatment and a more modest response to glutamate (Fig 2D). Moreover, while 18 interactions were individually significantly different (by ANC statistics) in comparisons of WT starved vs. stimulated, only 4 were significantly different in Shank3B^-/-^ comparisons (Table S2). These data, exemplified by mTOR_Raptor (Fig 2E), indicate that Shank3 neurons are less efficient at transmitting signals through the mTOR interaction network.

**Figure 2:**
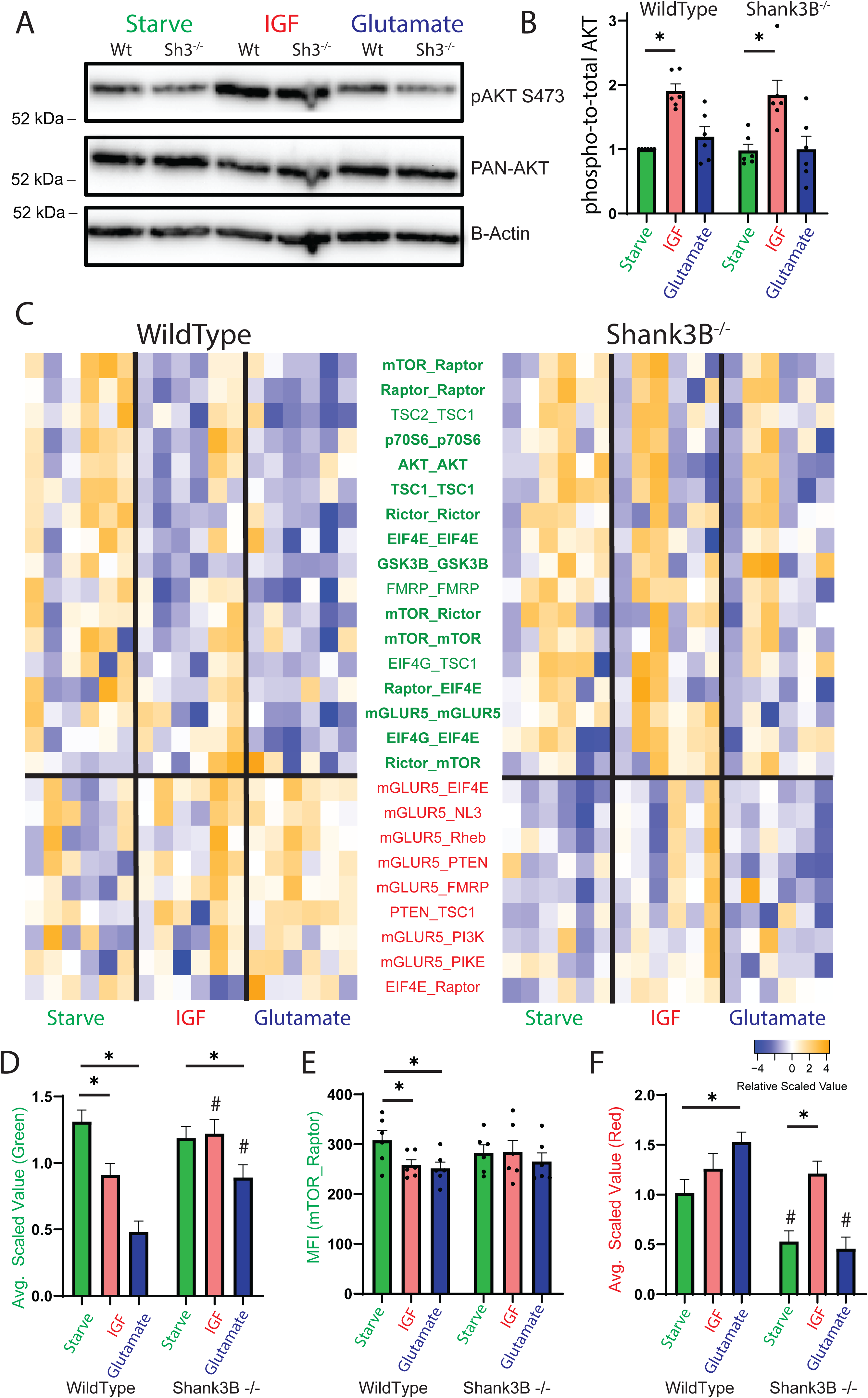
mTOR dynamics following IGF and Glutamate treatment of Shank3 KO neurons. A) Representative western blots showing phospho- and total AKT with actin for a loading control. B) Quantification of blots shown in A. * indicates p<0.05 by ANOVA followed by Dunnett post-hoc testing N =6. C) Heatmap of all significantly altered interactions as in Fig 1. D) Mean scaled value of all interactions in the green module. E) Median fluorescent intensity of the mTOR_Raptor co-association. F) Mean scaled value of all interactions in the red module. For D-F, * indicates p<0.05 compared to same genotype (treatment effect), # indicates p<0.05 compared to same treatment condition (genotype effect) by 2-way ANOVA followed by Tukey post-hoc testing (D,F) or ANC (E).

CNA also identified a “red” module that was significantly correlated with genotype (correlation coefficient = -0.41, p = 0.01), comprised largely of mGluR5-containing interactions (although note that mGluR5_mGluR5 was a member of the green/stimulation module). The average scaled value of the red/genotype module was decreased compared to WT in starved Shank3B^-/-^ neurons, and while Glutamate stimulation increased module intensity in WT neurons, it did not affect Shank3B^-/-^ neurons. Rather, only IGF stimulation increased red/genotype interactions in Shank3B^-/-^ neurons (Fig 2F). Dysregulation of the mGluR system in Shank3B^-/-^ mice is consistent with a known loss of mGluR scaffolding by Shank3 (*27*).

### mTOR activation following homeostatic up- or down-scaling

To study mTOR activation in a more physiologically relevant setting, we used the well-established model of in vitro homeostatic scaling (*29*) to induce synchronized synaptic plasticity in a population of neurons in vitro. DIV 18-20 mouse neurons were treated with tetrodotoxin (TTX) to induce upscaling or with Bicuculline (BIC) to induce downscaling and lysed after 12 or 48 hours. We first quantified mTOR signal transduction with phospho-western blots (Fig 3A). AKT phosphorylation was significantly but transiently reduced by TTX treatment at 12 hours, then significantly increased by BIC treatment at 48 hours (Fig 3B). Downstream of AKT, mTOR phosphorylation was reduced by TTX at 12 and 48 hours, but unchanged by BIC (Fig 3C). Phosphorylation of the mTORC1 substrate P70S6K1 was increased by BIC at 48 hours (Fig 3D), while phosphorylation of 4EBP1, another mTORC1, substrate was unchanged (Fig 3F). However, the total amount of 4EBP1 compared to Actin was reduced by TTX at 48 hours (Fig 3G).

**Figure 3:**
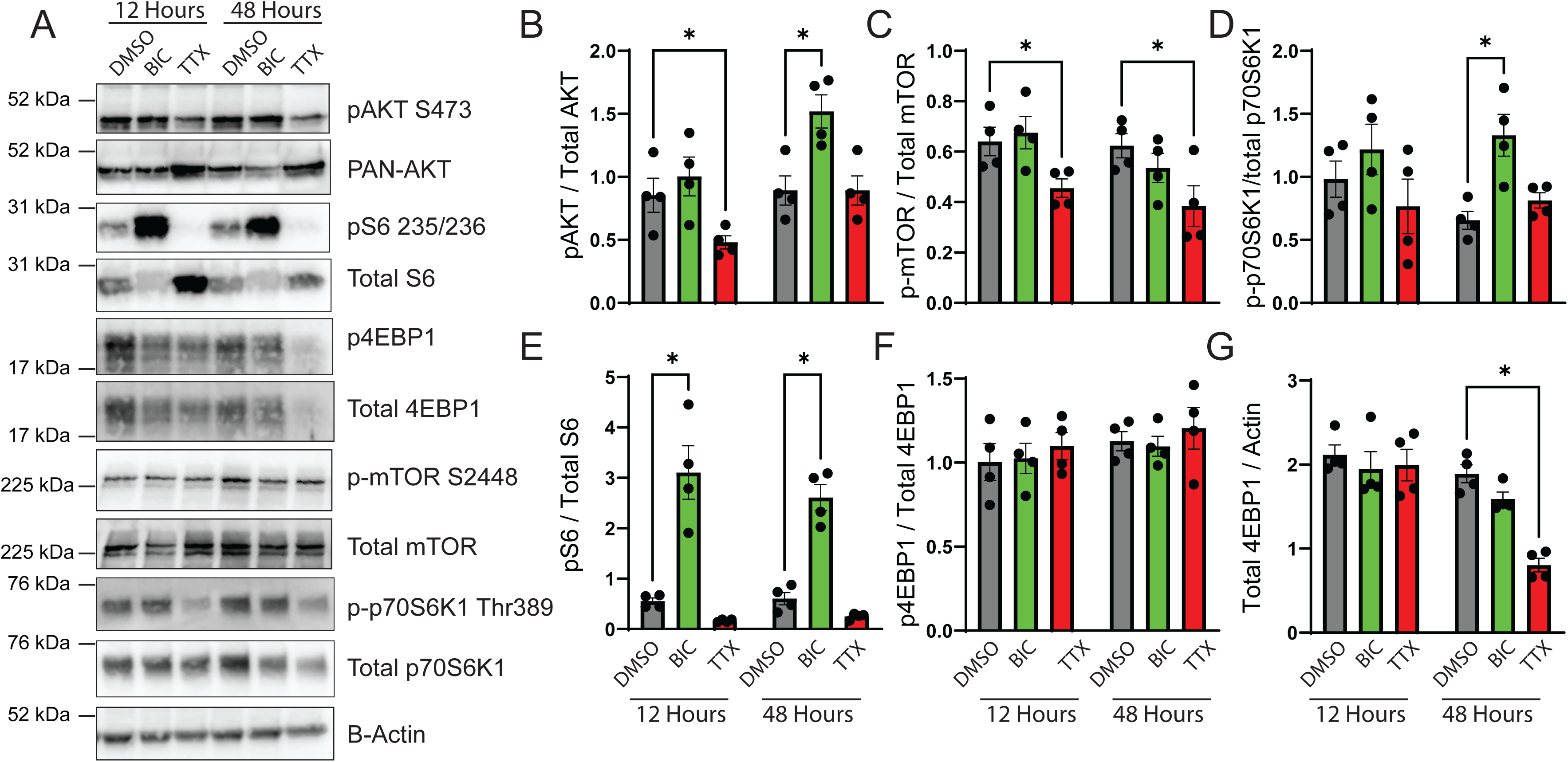
mTOR pathway phosphorylation following homeostatic scaling. A) Representative western blots showing phospho- and total AKT, S6, 4EPB1,mTOR and p70S6K following 12 or 48 hours of up- or down-scaling with TTX or BIC. B-G) Quantification of blots shown in A. * indicates p<0.05 compared to DMSO by 2-way ANOVA followed by Dunnett post-hoc testing, p<0.05. N =4.

Meanwhile, phosphorylation of P70S6K1’s substrate S6 was strongly and significantly increased by BIC at 12 and 48 hours and trended toward a reduction by TTX (Fig 3E). Overall, these data reveal a complex pattern of phosphorylation of mTOR pathway components by homeostatic scaling, varying with both time and direction (up or down) of scaling.

We next used QMI to observe changes in mTOR protein complex assembly following scaling. CNA analysis revealed two modules, a “turquoise” module that strongly correlated with BIC treatment (CC = -0.79, p = 2 x 10^-11^), and a “blue” module that best correlated with a hypothesis of activation in response to both treatments that increased over time (CC = -0.76, p = 5 x 10^-10^) (Fig 4A). The average scaled value of significant “turquoise module” interactions showed a reduction of module interactions at 12- and 48-hours following BIC, but no change with TTX at either timepoint (Fig 4B).

**Figure 4:**
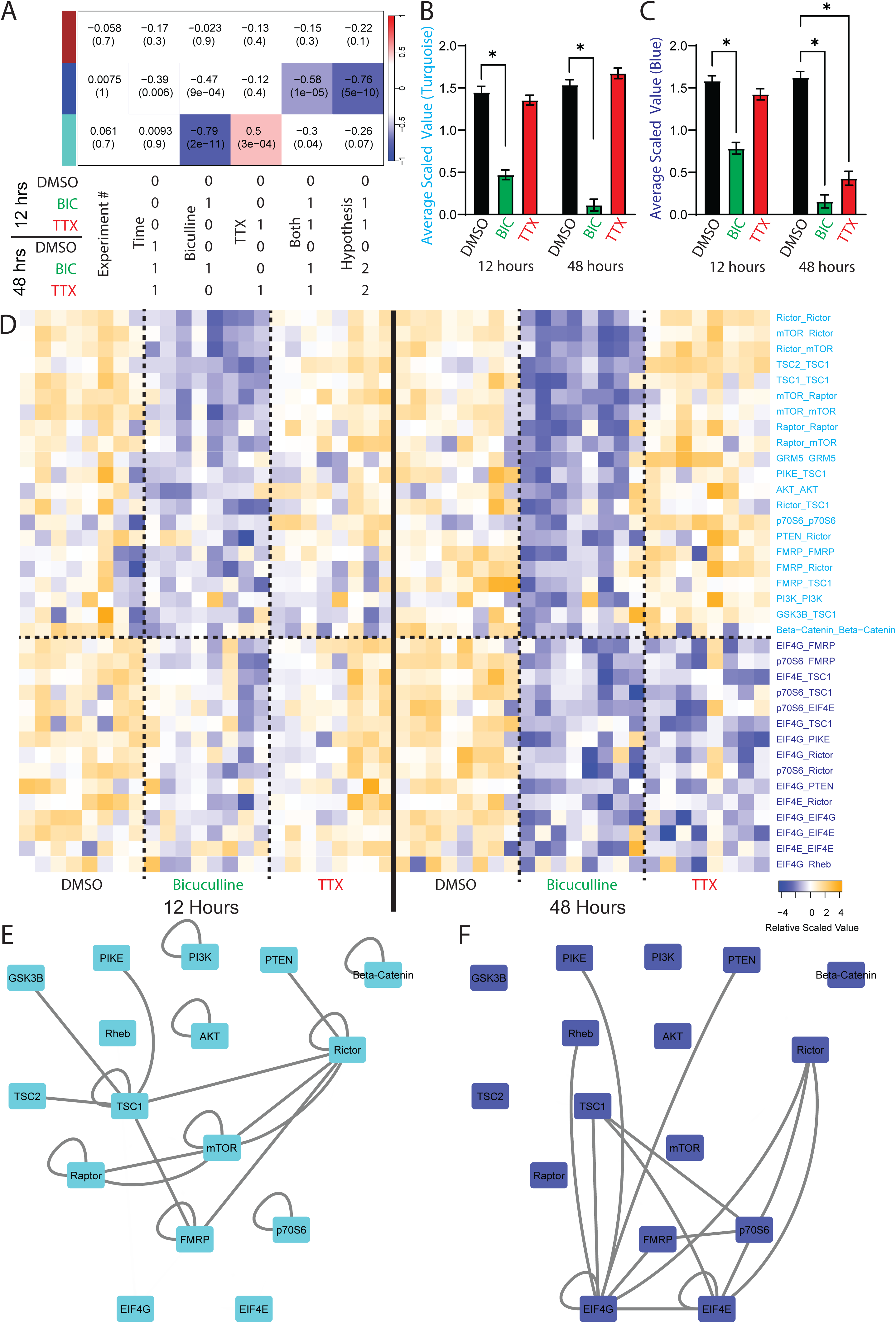
mTOR network dynamics following homeostatic scaling in cultured neurons. A) Module-trait table showing the correlation coefficient (top number) and p-value (bottom number) between the eigenvector of each color-coded module (colored rectangles on the left) and binary-coded trait labels shown in the table below. B,C) Average scaled value of all interactions in the turquoise (B) and blue (C) modules. * indicates p<0.05 by 2-way ANOVA followed by Dunnett post-hoc testing. D) Heatmap of the scaled values of all significantly altered interactions, N=8 per condition. Statistical significance calculated by ANC and CNA statistics. E,F) Node-edge diagrams of all interactions in the turquoise (E) and blue (F) modules. Nodes represent proteins measured by the QMI panel, edges represent interactions significantly by ANC and CNA statistics.

The “blue” module showed a reduction at 12 hours with BIC, and a larger reduction at 48 hours with both TTX and BIC treatment. Thus, the mTOR network differentiates between up- and down-scaling by slow activation of a single shared “Blue” module for upscaling vs. more rapid activation of the “blue” module <u>AND</u> activation of a unique BIC-only “turquoise” module for down-scaling.

We next examined the specific interactions that comprise each module (Fig 4D). Rictor_Rictor was the interaction most strongly correlated with the “Turquoise/BIC-only” module and was significantly reduced in 12 and 48-hour BIC treated neurons (Fig S2A).

Both mTORC1 (mTOR_Raptor (Fig S2B)) and TORC2 interactions were reduced in response to BIC. Three of 21 interactions in the BIC-only module were bi-directional: TSC2_TSC1 (Fig S2C), GRM5_GRM5 (Fig S2D) and PI3K_PI3K increased with TTX but decreased with BIC. Otherwise, the majority of interactions in the turquoise module were unchanged by TTX.

Interactions in the “blue/Both” module, exemplified by EIF4G_FMRP (Fig S2E) were significantly reduced by BIC at 12 hours, and further reduced by BIC and TTX at 48 hours. This pattern was consistent for the co-association between p70S6K_TSC1 (Fig S2F) and EIF4G_EIF4E. While the turquoise/BIC module involved more upstream members of the mTOR cascade including TORC2, TORC1 and the TSC complex (Fig 4E), members of the “Both/Blue” module reflected downstream proteins EIF4E, EIF4G and FMRP involved in translational output (Fig 4F). These data indicate that widespread activation of TORC1 and TORC2 is exclusive to BIC, but dissociation of EIF-containing translation complexes is shared between up- and down-scaling.

### Homeostatic scaling in Shank3B^-/-^ neurons

We next compared scaling in Shank3B^-/-^ neurons. Western blot for phospho-S6 showed up- and down-regulation in response to BIC and TTX, respectively, in both genotypes (Fig 5A) (although note that BIC was not significantly different from DMSO in Shank3B^-/-^ animals, (Fig S3A)). CNA identified two modules, “Brown” and “Turquoise”, that responded to BIC exclusively, and a single “Blue” module that responded to both treatments. The modules showed moderate consistency with those identified in the WT-only experiment (i.e. Figure 4), with 15/23 interactions in the two “BIC only” modules also appearing in the prior “BIC only” module, and 12/21 “Both” interactions appearing in the prior “Both” module. (Fig S3B,C). This partial overlap is a common feature of QMI data; due to the stringency of the statistical analysis, many interactions that are identified as “significant” in one experiment may only trend towards significance in the next (*24*). A heatmap of module interactions (Fig 5C) showed clear differences between Shank3B^-/-^ and WT signaling, which was more obvious when interactions were clustered by treatment (Fig S3D). In both genotypes, the turquoise “BIC only” module showed a strong reduction in response to BIC and a small increase in response to TTX. However, the average scaled value of the module was significantly lower in Shank3B^-/-^ animals, both at baseline and following TTX treatment (Fig 5D). For some interactions, such as Rictor_Rictor, Shank3B^-/-^ samples did not display a significant response to BIC treatment, while wildtype samples did (Fig 5E). For others such as TSC2_TSC1, there was still a significant effect of TTX and BIC treatment in Shank3B^-/-^, but the abundance of each interaction was significantly lower than the WT condition (Fig 5F). Similarly, for the “Brown” module, BIC treatment significantly reduced the averaged scaled value both WT and Shank3B^-/-^ neurons, but the magnitude of this reduction was greater in WT neurons (Fig 5G); the MFI of the most highly correlated interaction, PIKE_TSC1, was significantly reduced only in WT (Fig 5H). Overall, in the two “BIC-responsive” modules, signaling capacity was reduced in Shank3B^-/-^ neurons due to both a trend towards an activated state in untreated neurons (the Turquoise module), and a reduced magnitude of activation in response to BIC (the Brown module).

**Figure 5:**
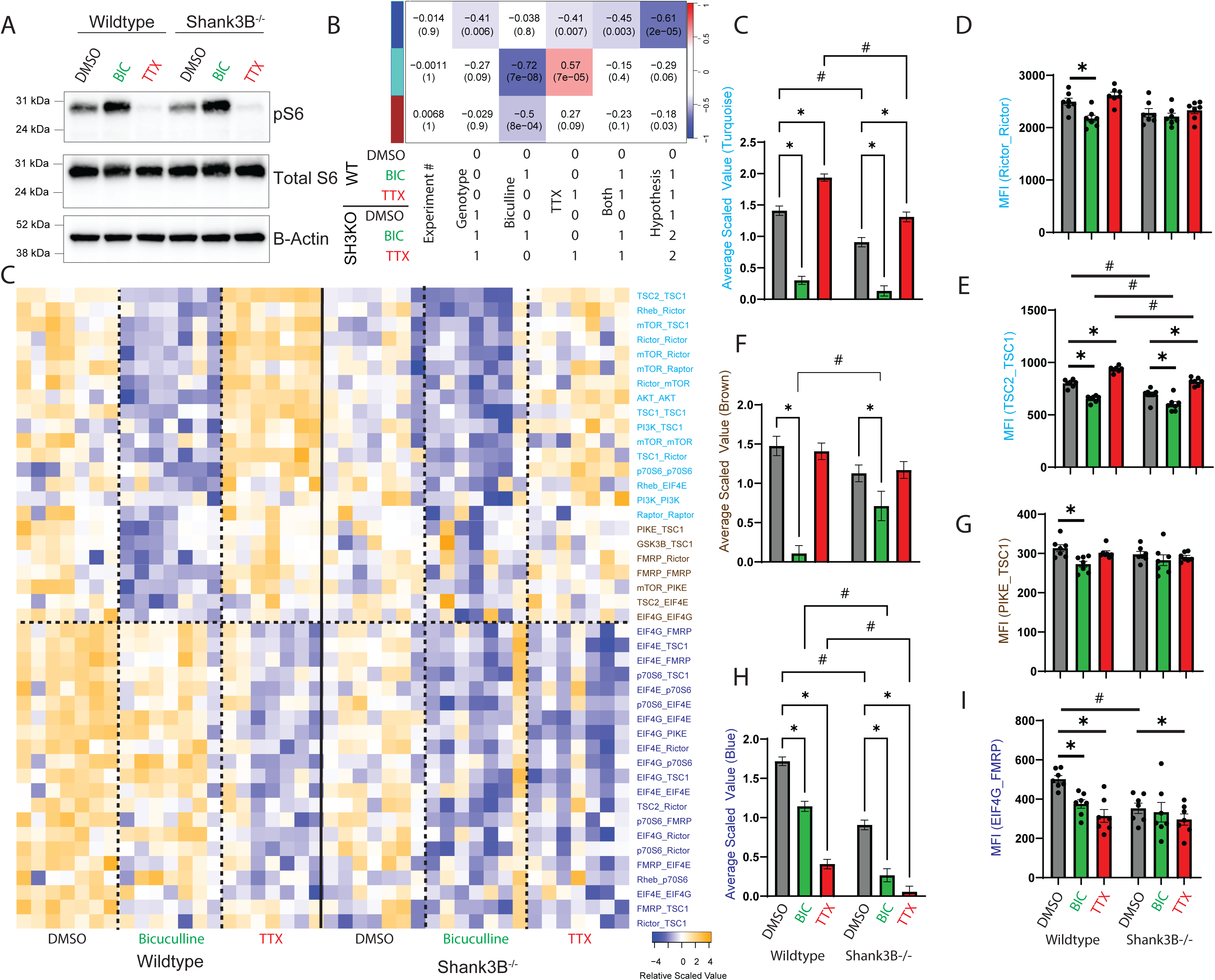
Homeostatic scaling in Shank3^-/-^ neurons. A) Western blot showing phospho- and total S6 following 48 hours of up- or down-scaling. B) Module-trait table showing the correlation coefficient (top number) and p-value (bottom number) between the eigenvector of each color-coded module (colored rectangles on the left) and binary-coded trait labels shown in the table below. C) Heatmap of the scaled values of all significantly altered interactions, by ANC and CNA, N=7 per condition. D-J) Mean scaled value of all interactions in each module (D,G,I), and representative interactions from each module (E,F,H,J). * indicates p<0.05 by 2-way ANOVA followed by Dunnett post-hoc testing (D,G,I) or ANC (F,E,H,J).

The BIC and TTX-responsive “Blue” module also showed significantly reduced averaged scaled intensity following TTX and BIC treatment in both genotypes (Fig 5I). Again, the “Blue” module’s average scaled value was significantly lower in Shank3B^-/-^ neurons compared to WT, in all conditions. The interaction most strongly correlated to the blue module, EIF4G_FMRP, was significantly reduced by BIC and TTX in WT neurons, but Shank3B^-/-^ neurons showed significantly reduced MFI in the DMSO condition compared to WT, and only a small non-significant response to BIC was observed (Fig 5J). Overall, the mTOR network appears “pre-activated” in unmanipulated Shank3B^-/-^ neurons because the constituent interactions are at levels more similar to the wildtype’s stimulated state than the wildtype’s unstimulated state. Functionally, this pre-activation reduces the dynamic range available to the network upon stimulation, rendering the system less sensitive to stimuli that rely on mTOR for signal transduction.

### Inhibition of mTOR signaling with Rapalink

Rapalink is a third generation mTOR inhibitor that is comprised of Rapamycin linked to the mTOR inhibitor Sapanisertib for maximal kinase inhibition of TORC1 through two independent mechanisms (*30*). Western blot indicated Rapalink-1 treatment inhibited phospho-S6 downstream of TORC1 without inhibiting phospho-AKT downstream of TORC2 in both WT and Shank3B^-/-^ neurons (Figure 6A-C). QMI analysis revealed two modules, turquoise that significantly correlated with Rapalink treatment (correlation coefficient = 0.84, p = 2*10^-7^), and “green” that correlated with both Rapalink (CC = -0.52, p = 0.009) and Genotype (CC=-0.43, p = 0.04). CNA is agnostic to the direction of change, but observation of the turquoise module heatmap (Fig 6D) revealed two classes of interactions—those that increased or decreased with Rapalink. Decreased interactions strongly and significantly reduced following Rapalink, and were not different between genotypes (Fig 6E). All decreased interactions involved mTOR itself, for example, mTOR_Rictor, reflecting conformational changes caused by Rapalink (Fig 6F). Critically, the remainder of the module strongly increased in abundance following Rapalink (Fig 6G), for example Rictor_TSC1 (Fig 6H). These data highlight the dissociative nature of the mTOR interaction network in neurons, since inhibition of a dissociative network should yield increased co-associations, as observed. The only interactions to decrease were mTOR-containing interactions, likely directly impacted by Rapalink binding.

**Figure 6:**
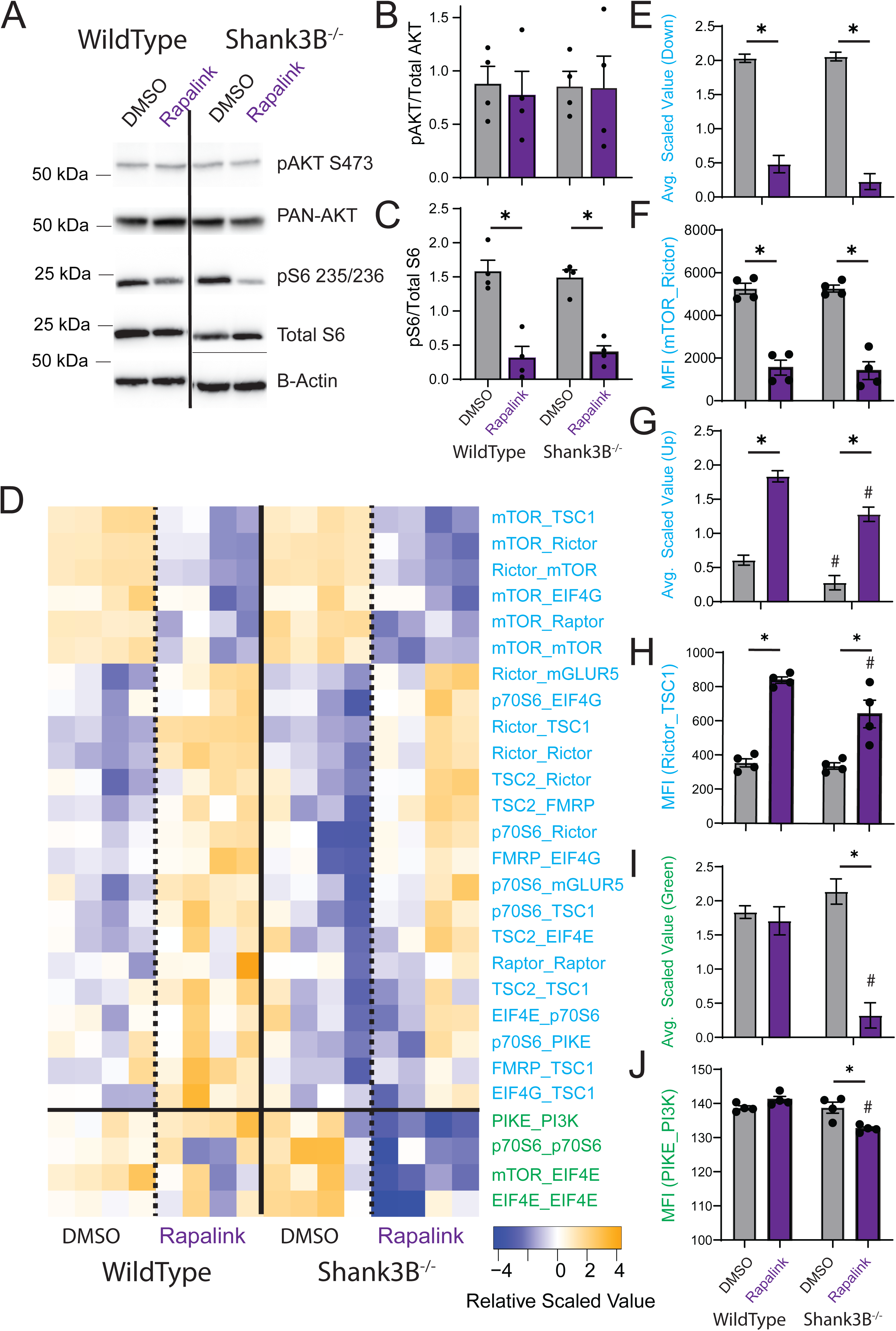
Inhibition of mTOR signaling with Rapalink in WT and Shank3B^-/-^ neurons. A) Representative western blots showing phospho- and total Akt, phospho- and total S6, and beta-actin in WT and Shank3B^-/-^ neurons with and without Rapalink treatment. B-C) Quantification of blots shown in A. * indicates p<0.05 by 2-way ANOVA followed by Dunnett post-hoc testing; N=4. D) Heatmap showing the scaled values of significantly different interactions identified by both ANC and CNA; N=4. E-J) Mean scaled value of all interactions in each module (E,G,I), and representative interactions from each module (F,H,J). * indicates p<0.05 compared to DMSO control within genotype (treatment effect), and # indicates p<0.05 compared to wildtype with same treatment (genotype effect) by 2-way ANOVA followed by Dunnett post-hoc testing (E,G,I) or ANC (F,H,J).

Compared to WT neurons, untreated Shank3B^-/-^ neurons showed significantly reduced co-associations among “up” proteins in the turquoise module at baseline, consistent with a “pre-activated” model (Fig 6G). Following Rapalink, average scaled intensity of the turquoise-up module was still reduced compared to WT (Fig 6G).

Moreover, the green module contained 4 interactions that reduced intensity in response to Rapalink exclusively in Shank3B^-/-^ neurons (Fig 6I), for example PIKE_PI3K (Fig 6J), which may reflect tonic mGluR activation associated with Shank3 loss (*31*). These data demonstrate that Shank3B^-/-^ deficient neurons have an abnormal dynamic range during inhibition with Rapalink, similar to the disrupted dynamic range observed during homeostatic scaling.

## Discussion

### Phosphorylation correlated with dissociation of interactions

The mTOR network has been extensively characterized over the past two decades (*32*), with the majority of studies conducted in actively proliferating or immortalized cell types, such as HEK293 and HeLa cells (*33*). Following stimulation, pathway components are typically phosphorylated according to a traditional linear hierarchy of PI3K -> AKT -> mTOR -> S6 kinase. Concurrent with phosphorylation, mTOR complexes assemble following stimulation; for example, Raptor-containing TORC1 complexes are recruited to the lysosome upon amino acid stimulation, which recruit TSC2 to the lysosome upon insulin stimulation to regulate Rheb activity (*34–36*). TORC2 complex assembly is enhanced upon growth factor signaling, andTORC2 localization to the plasma membrane is necessary for its role in phosphorylating AKT (*37*). Furthermore, interactions between TORC1 and ULK1 modulate autophagy initiation, demonstrating that mTOR complex dynamics extend beyond protein synthesis and cellular growth (*38*). Importantly, in immortalized cell types, mTOR phosphorylation and protein complex formation are positively correlated (*19*).

In neurons, mTOR is activated by stimuli such as brain-derived neurotrophic factor (BDNF) and synaptic activity, which lead to phosphorylation of pathway components like S6 and 4E-BP1. Several studies have utilized RAPTOR or RICTOR knockout mice (*17*, *39*) and Rapalogs (*40–43*) to demonstrate that both TORC1 and TORC2 are required for different types of synaptic plasticity. However, these studies have not considered mTOR complex dynamics. Meanwhile, some studies have made the counterintuitive observation that dendritic protein synthesis is reduced after acute mTOR activation (*44*, *45*), indicating that mTOR may function differently in neurons. However, direct evidence of stimulus-dependent mTOR complex assembly or disassembly in neurons remains limited.

Here we show that mTOR complexes disassemble following stimulation, despite *increases* in phosphorylation. These data imply that neurons have evolved a fundamentally different mechanism of mTOR regulation that reflects their unique usage of the system to mediate synaptic plasticity and dendritic excitability (*46*, *47*). Since mTOR controls local dendritic translation of proteins downstream of synaptic activity, we speculate that mTOR activation may down-regulate protein synthesis acutely following strong synaptic activity, giving a localized dendritic segment a temporary pause in translation. Such a pause would be consistent with imaging data showing a resistance to structural LTP (sLTP) following an initial sLTP-inducing stimulation (*48*), and the de-activation of NMDA receptors by GRM5 following activity-dependent untethering of GRM5 from its Shank-Homer scaffolding by phosphorylation or Homer1a expression (*27*, *49*, *50*). Together, these systems may allow a potentiated synapse to pause and process the incoming signal before being receptive to further potentiation. This hypothesis is also consistent with mTOR being a voltage sensor that regulates the excitability of dendritic segments (*51*).

### Combinatorial encoding of up- vs. down-scaling

Cells use a limited number of signal transduction pathways to perform a vast number of computations, begging the question, how can cells differentiate signals given such a limited set of pathways? Pawson postulated that different combinations of protein-protein interactions could encode different stimuli by activating combinations of signaling pathways (*2*). Indeed, we previously showed that a glutamate synapse protein interaction network recruits different combinations of protein interactions in response to NMDA vs. DHPG stimulation(*52*), consistent with the Pawson hypothesis. Alternatively, the intensity of activation of a conserved network could encode information. For example, in developing T cells, strong activation of a signaling network downstream of the T cell receptor encodes a cell death response to prevent self-recognition and autoimmunity, while weak activation of the same protein network encodes successful TCR recombination and permits continued development (*53*). Here, we identified a unique “A vs. A+B” encoding of competing stimuli. TTX-induced dissociation of a module containing downstream EIF-containing interactions, suggesting it induced a change in the translational output of the system. BIC induced dissociation of the same module, of the same approximate magnitude, to TTX. This dissociation is consistent with SILAC labeling mass spectrometry experiments that found large clusters of proteins that decreased synthesis in both TTX- and BIC-treated neurons (*54*). In addition, BIC induced dissociation of a second module not affected by TTX, consisting of more upstream interactions including TORC2 and the TSC complex. While both up- and down-scaling share common modulation and signaling changes through pathways such as the CaMKK/CaMKIV pathways, other signaling elements like brain-derived neurotrophic factor (BDNF), Arc, and PSD95 that are essential for up-scaling are dispensable for down-scaling (*55*). To our knowledge, this is the first time this type of A/A+B encoding has been documented.

### Homeostatic setpoint in Shank3B^-/-^ animals

Shank3 is an important scaffolding protein for the post-synaptic density, regulating the arrangement of its receptors and consequently their downstream signaling cascades(*27*, *28*, *56*, *57*). The loss of Shank3 causes hypoactive mTOR signaling by disrupting mGluR5 scaffolding, which both tonically activates the mGluR receptor in a ligand-independent manner, and impairs ligand-induced ERK and mTOR signaling downstream of the mGluR5 receptor (*56*, *58*). Consistent with ligand-independent activation, we observed reduced interactions in the unstimulated condition (Fig 5C,H, Fig 6G), corresponding to a partial or tonic activation of the dissociative mTOR interaction network. Moreover, we previously reported that synaptic signaling networks in Shank3B^-/-^ cortical neurons showed evidence of tonic hyperactivity. Protein interactions among a glutamate synapse interaction network dissociated upon plasticity-inducing stimulation, and were lower in resting Shank3B^-/-^ mutants, consistent with tonic activation of a protein network downstream of synaptic activity (*24*). Here, we find similar evidence that Shank3 loss mimics a partial activation of signal transduction in resting neurons, leading to reduced dynamic range upon signal activation.

However, the lack of response to IGF, and the blunted response to glutamate (Fig 2D,F) and homeostatic scaling (Fig 5F) suggest a more complex phenotype that mirrors the complex behavior of the mTOR network. For example, studies using acute shRNA-based knockdown of Shank3 (as in (*56*)) vs. genetic knockdown may lead to different findings due to extensive feedback among the mTOR network leading to homeostatic compensation and network re-arrangement, as is evident in experiments involving chronic vs. long-term treatment with Rapalogs (*59*). Further, the effects of Shank3 mutation can be developmentally regulated; for example cortico-striatal synapses are hyperactive in young Shank3 animals, but hypoactive in adults (*60*). Of note, Shank3 overexpression also causes mTOR hypoactivation and reduced mTOR phosphorylation (*61*). Future work should characterize the mechanisms of the Shank3’s regulation of the mTOR network to determine how it influences the response range of mTOR signaling and how this affects neurodevelopment. Elucidating the relationship between Shank3 and mTOR signaling will be critical in attempting to restore the setpoint of dysregulated signal transduction systems, which may be an avenue for normalizing system behavior.

## Materials and Methods

### Animals

All work with animals was performed in compliance with the Seattle Children’s Research Institute Institutional Animal Care and Use Committee under approved protocol no. 00072 and federal guidelines. CD1 and Shank3^tm2Gfng^ (stock 017688) mice were originally obtained from The Jackson Lab-oratory (Bar Harbor, ME) and maintained through in-house breeding. Mice were kept on a 12-12 light cycle with food and water available ad libitum.

### Genotyping

Crude DNA extract (0.3 ul) (Kapa Biosystems) from ear punch tissue was used for genotyping the Shank3 allele with the following primers: 5′-GAGACTGATCAGCGCAGTTG-3′, 5′-TGACATAATCGCTGGCAAAG-3′, and 5′-GCTATACGAAGT-TATGTCGACTAGG-3′ using standard polymerase chain reaction protocols.

### Cortical neuron culture and drug treatment

Primary cortical neuron cultures were prepared as described in previous works(*62*, *63*). Whole cortex from P0 or P1 mouse neonates was dissociated using papain (Worthington) and plated at a 1-1.2 x 10^6^ cell density per well in six well plates treated with poly-D-lysine. Cells were cultured in Neurobasal A medium supplemented with 2% B27 and 2 mM GlutaMAX (Thermo Fisher Scientific) and 100U/ml pennicillin and 100ug/ml streptomycin and kept at 37°, 5% CO_2_ for 17-21 days. After 3 to 5 days in vitro (DIV), 5-fluoro-2’-deoxyuridine was added to a final concentration of 5 µM to inhibit glial proliferation. For in vitro homeostatic scaling experiments, TTX (2 µM) or BIC (40 µM) (Tocris) was added directly to the culture medium, and cells were cultured as normal and then lysed after either 12 or 48 hours. DMSO (0.1%) served as a vehicle control.

Two days prior to mTOR stimulation and inhibition experiments, cultured neurons were given 680 µL of fresh media [400 µl] and pure water [280 µl] (to compensate for evaporation) to standardize the media nutrient composition. Rapalink (Cat# HY-111373 MedChem Express) was diluted in Neurobasal medium that lacked any supplements and added at 10nM to the culture medium. Neurons were cultured as normal and lysed after 3 hours. DMSO served as a vehicle control.

### Lysate Preparation

After drug treatment, neuron culture medium was removed, cells were washed twice with ice-cold dPBS, and cells were scraped in cold lysis buffer [150 mM NaCl, 50 mM Tris (pH 7.4), 1% Digitonin, 10 mM sodium fluoride, 2 mM sodium orthovanadate, protease inhibitor cocktail (Sigma-Aldrich), and phosphatase inhibitor cocktail (Sigma-Aldrich). Lysate was transferred to a centrifuge tube, incubated on ice for fifteen minutes, and centrifuged at 15000g for fifteen minutes to remove nuclear and cellular debris. The protein concentration of the supernatant was determined using a Bradford assay (Pierce).

### Western Blotting

Proteins were separated by SDS-polyacrylamide gel electrophoresis and transferred to a polyvinylidene difluoride membrane. Membranes were blocked in 4-5% milk in TBST [0.05 M tris (pH 7.2), 0.15 M NaCl, 0.1% Tween 20] for 1 hour at room temperature and incubated with primary antibody diluted according to the manufacture’s instructions overnight at 4°C or for 1 hour at room temperature. Primary antibodies were detected using species-specific horseradish peroxidase-conjugated secondary antibodies. Blots were developed using Femto Maximum Sensitivity Substrate (Pierce) and imaged using either a Protein Simple imaging system or Azur Biosystem 600.

### Antibodies

The following antibodies for the purpose of western blotting were acquired from Cell Signaling Technologies: Phospho-AKT Ser473 (catalog number 4060), PAN-AKT (#2920), Phospho-p44/42 MAPK Thr202/Tyr204 (#4370), p44/42 MAPK (#4695), S6 Ribosomal Protein (#2317), Phospho-S6 Ribosomal Protein Ser235/236 (#4858), Phospho-p70S6 Kinase Thr389 (#97596), p70S6 Kinase (#9202), Phospho-4EBP1 Thr37/46 (#2855), 4EBP1 (#9452). Antibodies against Beta-Actin were purchased from GeneTex (catalog number 109639).

### Quantitative Multiplex Immunoprecipitation

QMI was performed as described previously(*64*). A master mix containing each antibody-coupled Luminex bead was prepared and distributed to lysates normalized for protein concentration. Samples were incubated overnight at 4°C on a rotator. The following day, samples were washed in cold FlyP buffer [50 mM tris (pH7.4), 100 mM NaCl, 1% bovine serum albumin, and 0.02% sodium azide] and distributed into twice as many wells of a 96-well plate as there were probe antibodies for technical replicates.

Biotinylated probe antibodies were added, and the plate was incubated at 4°C with gentle agitation for one hour. The resulting complexes were washed three times with FlyP buffer on an automatic plate washer. The samples were then incubated for thirty minutes with streptavidin-phycoerythrin at 4°C with gentle agitation. Samples were washed three times again and resuspended in 120 ul of cold FlyP buffer and processed with a customized refrigerated Bio-Plex 200.

### Statistical Analysis

#### General

Statistical comparisons were tested by 1 or 2-way ANOVA as appropriate followed by multiple-comparison-correction by Dunnett (1-way) or Tukey (2-way) in the Prism software (GraphPad). ANOVA details are available in the ANOVA Table S3.

Comparisons between QMI bead distributions were made using ANC, an approach that takes into account inter-replicate reproducibility and bead distribution width while adjusting the alpha cutoff to achieve a type I error of 0.05, see for details(*26*, *64*).

Modules of Interactions that covaried with experimental conditions were identified using weighted correlation network analysis(*25*). Bead distributions were collapsed and the median fluorescent intensity (MFI) value was averaged across technical replicates for input into the WGCNA package for R. Interactions with an MFI less than 100 were removed as noise, and batch effects were corrected using COMBAT(*65*). Power values giving the approximation of scale-free topology were determined using soft thresholding with a power adjacency function, and modules were determined by the TOM matrix function in WGCNA. Modules whose eigenvectors were correlated with an experimental trait (P < 0.05) were of interest. Interactions whose probability of membership in a module of interest was (P < 0.05) were considered “hits”. Interactions that CNA “hits” and significantly different by for a given experimental condition were considered high confidence interactions affected in that condition. For Principal component analysis, post-COMBAT, log2 transformed MFI values were input into the prcomp function in R.

## Acknowledgements

We thank members of the SEPS, Page and Millen laboratories for helpful discussions. This work was supported by NIH grant MH113545 to SEPS.

## Author Contributions

DTW and SEPS conceived the study. DTW, EAB, VS, FH performed the experiments. DTW and SEPS analyzed the data and wrote the manuscript with contributions from all authors. SEPS obtained funding and corresponded with the journal.

## References

1. N. G. Ahn, The MAP kinase cascade. Discovery of a new signal transduction pathway. Mol Cell Biochem 127–128, 201–209 (1993).

2. T. Pawson, P. Nash, Assembly of cell regulatory systems through protein interaction domains. Science 300, 445–452 (2003).

3. B. A. Liu, B. W. Engelmann, P. D. Nash, The language of SH2 domain interactions defines phosphotyrosine-mediated signal transduction. FEBS Letters 586, 2597–2605 (2012).

4. Y. Ma, J. Nicolet, Specificity models in MAPK cascade signaling. FEBS Open Bio 13, 1177–1192 (2023).

5. H. Yang, D. G. Rudge, J. D. Koos, B. Vaidialingam, H. J. Yang, N. P. Pavletich, mTOR kinase structure, mechanism and regulation. Nature 497, 217–223 (2013).

6. J. O. Lipton, M. Sahin, The Neurology of mTOR. Neuron 84, 275–291 (2014).

7. L. H. Nguyen, A. Bordey, Convergent and Divergent Mechanisms of Epileptogenesis in mTORopathies. Front Neuroanat 15, 664695 (2021).

8. G. M. Mirzaa, C. D. Campbell, N. Solovieff, C. Goold, L. A. Jansen, S. Menon, A. E. Timms, V. Conti, J. D. Biag, C. Adams, E. A. Boyle, S. Collins, G. Ishak, S. Poliachik, K. M. Girisha, K. S. Yeung, B. H. Y. Chung, E. Rahikkala, S. A. Gunter, S. S. McDaniel, C. F. Macmurdo, J. A. Bernstein, B. Martin, R. Leary, S. Mahan, S. Liu, M. Weaver, M. Doerschner, S. Jhangiani, D. M. Muzny, E. Boerwinkle, R. A. Gibbs, J. R. Lupski, J. Shendure, R. P. Saneto, E. J. Novotny, C. J. Wilson, W. R. Sellers, M. Morrissey, R. F. Hevner, J. G. Ojemann, R. Guerrini, L. O. Murphy, W. Winckler, W. B. Dobyns, Association of MTOR Mutations With Developmental Brain Disorders, Including Megalencephaly, Focal Cortical Dysplasia, and Pigmentary Mosaicism. JAMA Neurol 73, 836–845 (2016).

9. K. M. Huber, E. Klann, M. Costa-Mattioli, R. S. Zukin, Dysregulation of Mammalian Target of Rapamycin Signaling in Mouse Models of Autism. J. Neurosci. 35, 13836–13842 (2015).

10. G. Tang, K. Gudsnuk, S.-H. Kuo, M. L. Cotrina, G. Rosoklija, A. Sosunov, M. S. Sonders, E. Kanter, C. Castagna, A. Yamamoto, Z. Yue, O. Arancio, B. S. Peterson, F. Champagne, A. J. Dwork, J. Goldman, D. Sulzer, Loss of mTOR-dependent macroautophagy causes autistic-like synaptic pruning deficits. Neuron 83, 1131–1143 (2014).

11. A. E. Clipperton-Allen, D. T. Page, Pten haploinsufficient mice show broad brain overgrowth but selective impairments in autism-relevant behavioral tests. Human Molecular Genetics 23, 3490– 3505 (2014).

12. D. Ehninger, S. Han, C. Shilyansky, Y. Zhou, W. Li, D. J. Kwiatkowski, V. Ramesh, A. J. Silva, Reversal of learning deficits in a Tsc2+/− mouse model of tuberous sclerosis. Nat Med 14, 843–848 (2008).

13. C. Nicolini, Y. Ahn, B. Michalski, J. M. Rho, M. Fahnestock, Decreased mTOR signaling pathway in human idiopathic autism and in rats exposed to valproic acid. Acta Neuropathol Commun 3, 3 (2015).

14. J. Zhang, J.-X. Zhang, Q.-L. Zhang, PI3K/AKT/mTOR-mediated autophagy in the development of autism spectrum disorder. Brain Res Bull 125, 152–158 (2016).

15. G. Y. Liu, D. M. Sabatini, mTOR at the nexus of nutrition, growth, ageing and disease. Nat Rev Mol Cell Biol 21, 183–203 (2020).

16. S.-M. Shen, C. Zhang, M.-K. Ge, S.-S. Dong, L. Xia, P. He, N. Zhang, Y. Ji, S. Yang, Y. Yu, J.-K. Zheng, J.-X. Yu, Q. Xia, G.-Q. Chen, PTENα and PTENβ promote carcinogenesis through WDR5 and H3K4 trimethylation. Nat Cell Biol 21, 1436–1448 (2019).

17. P. J. Zhu, C.-J. Chen, J. Mays, L. Stoica, M. Costa-Mattioli, mTORC2, but not mTORC1, is required for hippocampal mGluR-LTD and associated behaviors. Nat Neurosci 21, 799–802 (2018).

18. V. Karalis, H. S. Bateup, Current Approaches and Future Directions for the Treatment of mTORopathies. Dev Neurosci 43, 143–158 (2021).

19. D. T. Wehle, C. S. Bass, J. Sulc, G. Mirzaa, S. E. P. Smith, Protein interaction network analysis of mTOR signaling reveals modular organization. Journal of Biological Chemistry 299 (2023).

20. M. Zhu, Y.-C. Qin, C.-Q. Gao, H.-C. Yan, X.-G. Li, X.-Q. Wang, Extracellular Glutamate-Induced mTORC1 Activation via the IR/IRS/PI3K/Akt Pathway Enhances the Expansion of Porcine Intestinal Stem Cells. J Agric Food Chem 67, 9510–9521 (2019).

21. W. Ru, Y. Peng, L. Zhong, S.-J. Tang, A role of the mammalian target of rapamycin (mTOR) in glutamate-induced down-regulation of tuberous sclerosis complex proteins 2 (TSC2). J Mol Neurosci 47, 340–345 (2012).

22. O. Bibollet-Bahena, G. Almazan, IGF-1-stimulated protein synthesis in oligodendrocyte progenitors requires PI3K/mTOR/Akt and MEK/ERK pathways. J Neurochem 109, 1440–1451 (2009).

23. J. Gerhart, M. Kirschner, The theory of facilitated variation. Proc Natl Acad Sci U S A 104 Suppl 1, 8582–8589 (2007).

24. W. E. Heavner, J. D. Lautz, H. E. Speed, E. P. Gniffke, K. B. Immendorf, J. P. Welsh, N. A. Baertsch, S. E. P. Smith, Remodeling of the Homer-Shank interactome mediates homeostatic plasticity. Sci. Signal. 14, eabd7325 (2021).

25. P. Langfelder, S. Horvath, WGCNA: an R package for weighted correlation network analysis. BMC Bioinformatics 9, 559 (2008).

26. S. E. P. Smith, S. C. Neier, B. K. Reed, T. R. Davis, J. P. Sinnwell, J. E. Eckel-Passow, G. F. Sciallis, C. N. Wieland, R. R. Torgerson, D. Gil, C. Neuhauser, A. G. Schrum, Multiplex matrix network analysis of protein complexes in the human TCR signalosome. Science Signaling 9, rs7–rs7 (2016).

27. E. Moutin, S. Sakkaki, V. Compan, N. Bouquier, F. Giona, J. Areias, E. Goyet, A.-L. Hemonnot-Girard, V. Seube, B. Glasson, N. Benac, Y. Chastagnier, F. Raynaud, E. Audinat, L. Groc, T. Maurice, C. Sala, C. Verpelli, J. Perroy, Restoring glutamate receptosome dynamics at synapses rescues autism-like deficits in Shank3-deficient mice. Mol Psychiatry, 1–14 (2021).

28. J. Peça, C. Feliciano, J. T. Ting, W. Wang, M. F. Wells, T. N. Venkatraman, C. D. Lascola, Z. Fu, G. Feng, Shank3 mutant mice display autistic-like behaviours and striatal dysfunction. Nature 472, 437–442 (2011).

29. G. G. Turrigiano, K. R. Leslie, N. S. Desai, L. C. Rutherford, S. B. Nelson, Activity-dependent scaling of quantal amplitude in neocortical neurons. Nature 391, 892–896 (1998).

30. V. S. Rodrik-Outmezguine, M. Okaniwa, Z. Yao, C. J. Novotny, C. McWhirter, A. Banaji, H. Won, W. Wong, M. Berger, E. de Stanchina, D. G. Barratt, S. Cosulich, T. Klinowska, N. Rosen, K. M. Shokat, Overcoming mTOR resistance mutations with a new-generation mTOR inhibitor. Nature 534, 272– 276 (2016).

31. C. Gross, C.-W. Chang, S. M. Kelly, A. Bhattacharya, S. M. J. McBride, S. W. Danielson, M. Q. Jiang, C. B. Chan, K. Ye, J. R. Gibson, E. Klann, T. A. Jongens, K. H. Moberg, K. M. Huber, G. J. Bassell, Increased Expression of the PI3K Enhancer PIKE Mediates Deficits in Synaptic Plasticity and Behavior in Fragile X Syndrome. Cell Reports 11, 727–736 (2015).

32. D. M. Sabatini, Twenty-five years of mTOR: Uncovering the link from nutrients to growth. PNAS 114, 11818–11825 (2017).

33. M. Laplante, D. M. Sabatini, mTOR signaling in growth control and disease. Cell 149, 274–293 (2012).

34. Y. Sancak, L. Bar-Peled, R. Zoncu, A. L. Markhard, S. Nada, D. M. Sabatini, Ragulator-Rag complex targets mTORC1 to the lysosomal surface and is necessary for its activation by amino acids. Cell 141, 290–303 (2010).

35. D.-H. Kim, D. D. Sarbassov, S. M. Ali, J. E. King, R. R. Latek, H. Erdjument-Bromage, P. Tempst, D. M. Sabatini, mTOR Interacts with Raptor to Form a Nutrient-Sensitive Complex that Signals to the Cell Growth Machinery. Cell 110, 163–175 (2002).

36. D. Meng, Q. Yang, M.-H. Jeong, A. Curukovic, S. Tiwary, C. H. Melick, T. D. Lama-Sherpa, H. Wang, M. Huerta-Rosario, G. Urquhart, L. G. Zacharias, C. Lewis, R. J. DeBerardinis, J. L. Jewell, SNAT7 regulates mTORC1 via macropinocytosis. Proc Natl Acad Sci U S A 119, e2123261119 (2022).

37. D. D. Sarbassov, S. M. Ali, D.-H. Kim, D. A. Guertin, R. R. Latek, H. Erdjument-Bromage, P. Tempst, D. M. Sabatini, Rictor, a novel binding partner of mTOR, defines a rapamycin-insensitive and raptor-independent pathway that regulates the cytoskeleton. Curr Biol 14, 1296–1302 (2004).

38. J. Kim, M. Kundu, B. Viollet, K.-L. Guan, AMPK and mTOR regulate autophagy through direct phosphorylation of Ulk1. Nat Cell Biol 13, 132–141 (2011).

39. M. P. McCabe, E. R. Cullen, C. M. Barrows, A. N. Shore, K. I. Tooke, K. A. Laprade, J. M. Stafford, M. C. Weston, Genetic inactivation of mTORC1 or mTORC2 in neurons reveals distinct functions in glutamatergic synaptic transmission. eLife 9, e51440 (2020).

40. M. Pereyra, C. Katche, A. B. de Landeta, J. H. Medina, mTORC1 controls long-term memory retrieval. Sci Rep 8, 8759 (2018).

41. F. E. Henry, X. Wang, D. Serrano, A. S. Perez, C. J. L. Carruthers, E. L. Stuenkel, M. A. Sutton, A Unique Homeostatic Signaling Pathway Links Synaptic Inactivity to Postsynaptic mTORC1. J Neurosci 38, 2207–2225 (2018).

42. S. J. Tang, G. Reis, H. Kang, A.-C. Gingras, N. Sonenberg, E. M. Schuman, A rapamycin-sensitive signaling pathway contributes to long-term synaptic plasticity in the hippocampus. Proc Natl Acad Sci U S A 99, 467–472 (2002).

43. L. Stoica, P. J. Zhu, W. Huang, H. Zhou, S. C. Kozma, M. Costa-Mattioli, Selective pharmacogenetic inhibition of mammalian target of Rapamycin complex I (mTORC1) blocks long-term synaptic plasticity and memory storage. Proc Natl Acad Sci U S A 108, 3791–3796 (2011).

44. K. F. Raab-Graham, P. C. G. Haddick, Y. N. Jan, L. Y. Jan, Activity- and mTOR-dependent suppression of Kv1.1 channel mRNA translation in dendrites. Science 314, 144–148 (2006).

45. F. Niere, S. Namjoshi, E. Song, G. A. Dilly, G. Schoenhard, B. V. Zemelman, Y. Mechref, K. F. Raab-Graham, Analysis of Proteins That Rapidly Change Upon Mechanistic/Mammalian Target of Rapamycin Complex 1 (mTORC1) Repression Identifies Parkinson Protein 7 (PARK7) as a Novel Protein Aberrantly Expressed in Tuberous Sclerosis Complex (TSC) *. Molecular & Cellular Proteomics 15, 412–430 (2016).

46. S. A. Swanger, G. J. Bassell, Dendritic protein synthesis in the normal and diseased brain. Neuroscience 232, 106–127 (2013).

47. A.-S. Hafner, P. G. Donlin-Asp, B. Leitch, E. Herzog, E. M. Schuman, Local protein synthesis is a ubiquitous feature of neuronal pre- and postsynaptic compartments. Science 364, eaau3644 (2019).

48. J. C. Flores, D. Sarkar, K. Zito, A synapse-specific refractory period for plasticity at individual dendritic spines. Proceedings of the National Academy of Sciences 122, e2410433122 (2025).

49. J. Perroy, F. Raynaud, V. Homburger, M.-C. Rousset, L. Telley, J. Bockaert, L. Fagni, Direct interaction enables cross-talk between ionotropic and group I metabotropic glutamate receptors. J Biol Chem 283, 6799–6805 (2008).

50. E. Moutin, F. Raynaud, J. Roger, E. Pellegrino, V. Homburger, F. Bertaso, V. Ollendorff, J. Bockaert, L. Fagni, J. Perroy, Dynamic remodeling of scaffold interactions in dendritic spines controls synaptic excitability. J Cell Biol 198, 251–263 (2012).

51. F. Niere, K. F. Raab-Graham, mTORC1 Is a Local, Postsynaptic Voltage Sensor Regulated by Positive and Negative Feedback Pathways. Front Cell Neurosci 11, 152 (2017).

52. J. D. Lautz, E. A. Brown, A. A. Williams VanSchoiack, S. E. P. Smith, Synaptic activity induces input-specific rearrangements in a targeted synaptic protein interaction network. Journal of Neurochemistry 146, 540–559 (2018).

53. S. C. Neier, A. Ferrer, K. M. Wilton, S. E. P. Smith, A. M. H. Kelcher, K. D. Pavelko, J. M. Canfield, T. R. Davis, R. J. Stiles, Z. Chen, J. McCluskey, S. R. Burrows, J. Rossjohn, D. M. Hebrink, E. M. Carmona, A. H. Limper, D. J. Kappes, P. J. Wettstein, A. J. Johnson, L. R. Pease, M. A. Daniels, C. Neuhauser, D. Gil, A. G. Schrum, The early proximal αβ TCR signalosome specifies thymic selection outcome through a quantitative protein interaction network. Sci Immunol 4, eaal2201 (2019).

54. A. R. Dörrbaum, B. Alvarez-Castelao, B. Nassim-Assir, J. D. Langer, E. M. Schuman, Proteome dynamics during homeostatic scaling in cultured neurons. eLife 9, e52939 (2020).

55. G. Turrigiano, Homeostatic synaptic plasticity: local and global mechanisms for stabilizing neuronal function. Cold Spring Harb Perspect Biol 4, a005736 (2012).

56. M. Bidinosti, P. Botta, S. Krüttner, C. C. Proenca, N. Stoehr, M. Bernhard, I. Fruh, M. Mueller, D. Bonenfant, H. Voshol, W. Carbone, S. J. Neal, S. M. McTighe, G. Roma, R. E. Dolmetsch, J. A. Porter, P. Caroni, T. Bouwmeester, A. Lüthi, I. Galimberti, CLK2 inhibition ameliorates autistic features associated with SHANK3 deficiency. Science 351, 1199–1203 (2016).

57. P. Monteiro, G. Feng, SHANK proteins: roles at the synapse and in autism spectrum disorder. Nat Rev Neurosci 18, 147–157 (2017).

58. A. Mossa, J. Pagano, L. Ponzoni, A. Tozzi, E. Vezzoli, M. Sciaccaluga, C. Costa, S. Beretta, M. Francolini, M. Sala, P. Calabresi, T. M. Boeckers, C. Sala, C. Verpelli, Developmental impaired Akt signaling in the Shank1 and Shank3 double knock-out mice. Mol Psychiatry 26, 1928–1944 (2021).

59. J. Li, S. G. Kim, J. Blenis, Rapamycin: one drug, many effects. Cell Metab 19, 373–379 (2014).

60. R. T. Peixoto, W. Wang, D. M. Croney, Y. Kozorovitskiy, B. L. Sabatini, Early hyperactivity and precocious maturation of corticostriatal circuits in Shank3B(-/-) mice. Nat. Neurosci. 19, 716–724 (2016).

61. Y. Lee, S. G. Kim, B. Lee, Y. Zhang, Y. Kim, S. Kim, E. Kim, H. Kang, K. Han, Striatal Transcriptome and Interactome Analysis of Shank3-overexpressing Mice Reveals the Connectivity between Shank3 and mTORC1 Signaling. Front. Mol. Neurosci. 10 (2017).

62. J. D. Lautz, K. B. Tsegay, Z. Zhu, E. P. Gniffke, J. P. Welsh, S. E. P. Smith, Synaptic protein interaction networks encode experience by assuming stimulus-specific and brain-region-specific states. Cell Rep 37, 110076 (2021).

63. L. G. W. Hilgenberg, M. A. Smith, Preparation of Dissociated Mouse Cortical Neuron Cultures. JoVE (Journal of Visualized Experiments), e562 (2007).

64. E. A. Brown, S. C. Neier, C. Neuhauser, A. G. Schrum, S. E. P. Smith, Quantification of Protein Interaction Network Dynamics using Multiplexed Co-Immunoprecipitation. JoVE (Journal of Visualized Experiments), e60029 (2019).

65. J. T. Leek, W. E. Johnson, H. S. Parker, A. E. Jaffe, J. D. Storey, The sva package for removing batch effects and other unwanted variation in high-throughput experiments. Bioinformatics 28, 882–883 (2012).

