## Supplement for "Dissociation of the mTOR protein interaction network following neuronal activation is altered by Shank3 mutation"

**Contents:**

**Table S1.**

**Table S2**

**Table S3**

**Figure S1**

**Figure S2**

**Table S1: Antibody pairs used for QMI**

| | Target | Uniprot identifier | Uniprot # | IP antibody clone | IP antibody Catalog number | IP antibody Supplier | Probe antibody clone | Probe antibody Catalog number | Probe antibody Supplier | Mouse Validation |
|---|---|---|---|---|---|---|---|---|---|---|
| 1 | AKT1 | AKT1_MOUSE | P31750 | SKB1 | 05-591 | Sigma Aldrich | B1 | sc-5298 | Santa Cruz Biotechnology | siRNA (Wehle et al 2023) |
| 2 | Beta-Catenin | CTNB1_MOUSE | Q02248 | 196624 | MAB13292 | R&D Systems | OTI2H7 | CF502409 | Thermo Fisher Scientific | Mouse KO liver lysate (B6.129-Ctnnb1tm2Kem/KnwJ) (Wehle et al 2023) |
| 3 | eIF4E | IF4E_MOUSE | P63073 | 299910 | MAB3228 | R&D Systems | Polyclonal | A301-153A | Fortis Life Sciences | siRNA (Wehle et al 2023) |
| 4 | EIF4G | IF4G1_MOUSE | Q6NZJ6 | G6899 | MA5-14971 | Thermo Fisher Scientific | 3A10 | SAB1403762 | Sigma-Aldrich | siRNA (Wehle et al 2023) |
| 5 | FMRP | FMR1_MOUSE | P35922 | Polyclonal | F4055 | Sigma Aldrich | 5C2 | 834701 | BioLegend | Mouse KO brain lystate (B6.129P2-Fmr1tm1Cgr/J) (Wehle et al 2023) |
| 6 | GSK3B | GSK3B_MOUSE | Q9WV60 | 3D6B4 | NBP2-52478 | Novus Biologicals | 3D10 | NBP1-47470 | Novus Biologicals | Mouse Embryonic fibroblast lysate (GSK3β(-/-); GSK3α(flox/-) (3/4KO) (Wehle et al 2023) |
| 7 | mGluR5 | GRM5_MOUSE | Q3UVX5 | 5675 | AB5675 | Sigma Aldrich | N75/3 | 75-115 | Antibodies Incorporated | Plasmid overexpression (Lautz et al 2018) |
| 8 | mTOR | MTOR_MOUSE | Q9JLN9 | 2ID8.2 | 05-1592 | Sigma Aldrich | 30 | sc-517464 | Santa Cruz Biotechnology | siRNA (Wehle et al 2023) |
| 9 | p70S6K | KS6A1_MOUSE | P18653 | 215247 | MAB8962 | R&D Systems | Polyclonal | AF8962 | Novus Biologicals | Mouse KO liver lysate (C.129(B6)-Rps6kb1tm1Gtho/NsoJ) (Wehle et al 2023) |
| 10 | PI3K | P85A_MOUSE | P26450 | U5 | MA1-74183 | Thermo Fisher Scientific | AB6 | 05-212 | Sigma Aldrich | Plasmid overexpression (Lautz et al 2018) |
| 11 | PIKE | AGAP2_MOUSE | Q3UHD9 | Polyclonal | A304-263A | Fortis Life Sciences | Polyclonal | 200-401-DN8 | Rockland | X63 Myeloma (Smith et al, 2016) |
| 12 | PTEN | PTEN_MOUSE | O08586 | 217702 | MAB847 | Novus Biologicals | 6H2.1 | 04-035 | Sigma Aldrich | Plasmid overexpression (Wehle et al 2023) |
| 13 | RAPTOR | RPTOR_MOUSE | Q8K4Q0 | Polyclonal | 42-4000 | Thermo Fisher Scientific | 514208 | MAB5957 | Novus Biologicals | siRNA (Wehle et al 2023) |
| 14 | Rheb | RHEB_MOUSE | Q921J2 | 344912 | MAB3426 | R&D Systems | JG37-12 | ET7108-44 | HUABIO | Plasmid overexpression (Wehle et al 2023) |
| 15 | RICTOR | RICTR_mouse | Q6QI06 | 7B3 | NBP1-51645 | Novus Biologicals | 9F1.2 | 05-1471 | Sigma Aldrich | mouse KO ESC lysate (Wehle et al 2023) |
| 16 | TSC1 | TSC1_MOUSE | Q9EP53 | 5C8A12 | 37-0400 | Thermo Fisher Scientific | Polyclonal | NB100-2315 | Novus Biologicals | Mouse Embryonic fibroblast lysate (TSC1(-/-)) (Wehle et al 2023) |
| 17 | TSC2 | TSC2_MOUSE | Q61037 | 614204 | MAB40401 | R&D Systems | SC05-59 | ET1610-10 | HUABIO | Plasmid overexpression (Wehle et al 2023) |

**Table S2: ANC results from WT and Shank3 IGF/Glutamate treatment**

| | WT | | MU | |
|---|---|---|---|---|
| | IGF v STARVE | GLUT vs STARVE | IGF v STARVE | GLUT vs STARVE |
| TSC1_TSC1 | -0.49 | -0.36 | | |
| GSK3B_GSK3B | -0.42 | | | |
| Raptor_Raptor | -0.37 | -0.44 | | |
| Rictor_Rictor | -0.34 | -0.28 | | |
| mTOR_Raptor | -0.30 | -0.23 | | |
| FMRP_FMRP | -0.27 | -0.55 | | -0.26 |
| EIF4E_EIF4E | -0.21 | | | |
| TSC2_TSC1 | -0.19 | -0.16 | | |
| mGLUR5_EIF4E | 0.29 | 0.25 | | |
| EIF4G_EIF4E | | -0.48 | | -0.51 |
| EIF4G_TSC1 | | -0.41 | | -0.24 |
| mGLUR5_mGLUR5 | | -0.33 | 0.04 | -0.29 |
| p70S6_p70S6 | | -0.32 | | |
| mTOR_mTOR | | -0.26 | | |
| AKT_AKT | | -0.26 | | |
| mTOR_Rictor | | -0.24 | | |
| mGLUR5_PI3K | | 0.02 | | |
| mGLUR5_Rheb | | 0.14 | | |
| Rictor_mTOR | | | 0.09 | |
| mGLUR5_Raptor | | | 0.21 | |
| Raptor_EIF4E | | | | -0.11 |

Numbers represent the log2 fold change shown only for ANC-significant interactions.

**Table S3: ANOVA Table for Figures 1-6**

| Figure | type | DFn, DFd | F | p value |
|---|---|---|---|---|
| 1C | 1-way | 3,12 | 24.75 | <0.0001 |
| 1H | 1-way | 3, 316 | 119.3 | <0.0001 |
| 1I | 1-way | 3, 12 | 7.366 | 0.0047 |
| 1J | 1-way | 3, 92 | 24.75 | <0.0001 |
| 1K | 1-way | 3, 12 | 3.032 | 0.071 |
| 1L | 1-way | 3, 60 | 3.022 | 0.0365 |
| 1M | 1-way | 3, 12 | 1.941 | 0.17 |
| 2B | 1-way | 5, 30 | 8.171 | <0.0001 |
| 2D | 2-way | treatment 2, 606 | 23.41 | <0.0001 |
| | | genotype 1, 606 | 12.93 | 0.0003 |
| | | interaction 2, 606 | 5.522 | 0.42 |
| 2F | 2-way | treatment 2, 318 | 6.921 | 0.0011 |
| | | genotype 1, 318 | 27.78 | <0.0001 |
| | | interaction 2, 318 | 8.401 | 0.0003 |
| 3B | 2-way | treatment 2, 18 | 20.51 | <0.0001 |
| | | time 1, 18 | 27.78 | 0.0302 |
| | | interaction 2, 18 | 8.401 | 0.1032 |
| 3C | 2-way | treatment 2, 18 | 7.711 | 0.0038 |
| | | time 1, 18 | 2.501 | 0.1312 |
| | | interaction 2, 18 | 0.5484 | 0.5872 |
| 3D | 2-way | treatment 2, 18 | 6.144 | 0.0092 |
| | | time 1, 18 | 0.1923 | 0.6662 |
| | | interaction 2, 18 | 1.176 | 0.331 |
| 3E | 2-way | treatment 2, 18 | 66.96 | <0.0001 |
| | | time 1, 18 | 0.3493 | 0.5618 |
| | | interaction 2, 18 | 0.873 | 0.4347 |
| 3F | 2-way | treatment 2, 18 | 0.6388 | 0.5395 |
| | | time 1, 18 | 1.876 | 0.1876 |
| | | interaction 2, 18 | 0.0435 | 0.9575 |
| 3G | 2-way | treatment 2, 18 | 9.491 | 0.0015 |
| | | time 1, 18 | 26.68 | <0.0001 |
| | | interaction 2, 18 | 6.982 | 0.0057 |
| 4B | 2-way | treatment 2, 954 | 246.1 | <0.0001 |
| | | time 1, 954 | 0.083 | <0.0001 |
| | | interaction 2, 954 | 14.81 | 0.7725 |
| 4C | 2-way | treatment 2, 762 | 127.4 | <0.0001 |
| | | time 1, 762 | 82.04 | <0.0001 |
| | | interaction 2, 762 | 27.15 | <0.0001 |
| 5C | 2-way | treatment 2, 666 | 197.7 | <0.0001 |
| | | genotype 1, 666 | 53.69 | <0.0001 |
| | | interaction 2, 666 | 5.365 | 0.0049 |
| 5F | 2-way | treatment 2, 288 | 33.22 | <0.0001 |
| | | genotype 1, 288 | 0.003 | 0.9531 |
| | | interaction 2, 288 | 8.617 | 0.0002 |
| 5H | 2-way | treatment 2, 876 | 131.3 | <0.0001 |
| | | genotype 1, 876 | 155.5 | <0.0001 |
| | | interaction 2, 876 | 9.157 | 0.0001 |
| 6B | 2-way | treatment 1, 12 | 0.078 | 0.865 |
| | | genotype 1, 12 | 0.009 | 0.923 |
| | | interaction 1, 12 | 0.044 | 0.783 |
| 6C | 2-way | treatment 1, 12 | 76.81 | <0.0001 |
| | | genotype 1, 12 | 0.0028 | 0.958 |
| | | interaction 1, 12 | 0.395 | 0.541 |
| 6E | 2-way | treatment 1, 268 | 305.9 | <0.0001 |
| | | genotype 1, 268 | 1.42 | 0.236 |
| | | interaction 1, 268 | 2.11 | 0.149 |
| 6F | | treatment 1, 12 | 147.9 | <0.0001 |
| | | genotype 1, 12 | 0.048 | 0.829 |
| | | interaction 1, 12 | 0.051 | 0.824 |
| 6G | 2-way | treatment 1, 268 | 145.1 | <0.0001 |
| | | genotype 1, 268 | 22.91 | <0.0001 |
| | | interaction 1, 268 | 1.48 | 0.224 |
| 6H | | treatment 1, 12 | 80.44 | <0.0001 |
| | | genotype 1, 12 | 5.76 | 0.033 |
| | | interaction 1, 12 | 4.17 | 0.063 |
| 6I | 2-way | treatment 1, 60 | 31.41 | <0.0001 |
| | | genotype 1, 60 | 9.78 | 0.0027 |
| | | interaction 1, 60 | 23.68 | <0.0001 |
| 6J | | treatment 1, 12 | 3.459 | 0.087 |
| | | genotype 1, 12 | 17.93 | 0.001 |
| | | interaction 1, 12 | 18.15 | 0.001 |

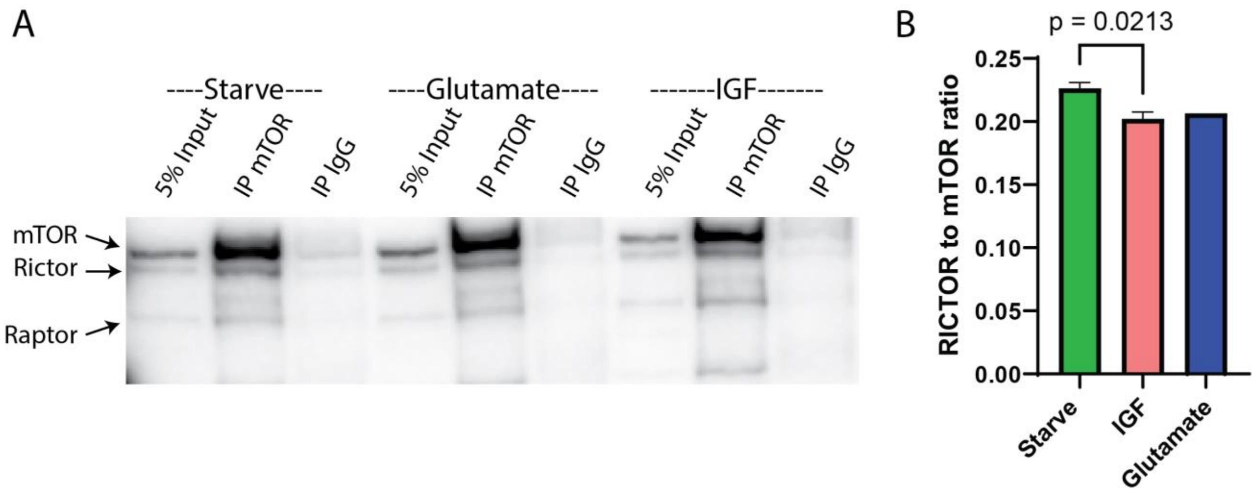


**Figure S1: IP-western blot of mTOR_Rictor.** Compare to QMI data in Fig 1I. A) Representative western blot B) Quantification of N=2 independent experiments. * indicates p<0.05 compared to the DMSO condition by ANC

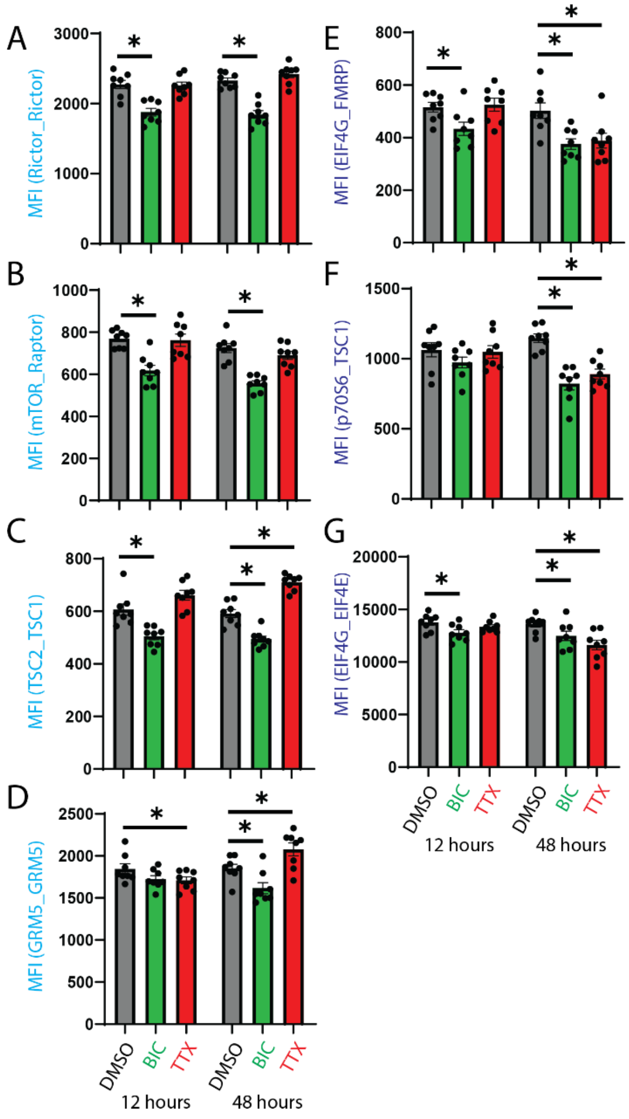


**Figure S2: Select interactions that respond to homeostatic upscaling. Related to Figure 4.** A-G) Median fluorescent intensity (MFI) for interactions in the turquoise (BIC) and blue (Both) modules. * indicates p<0.05 compared to the DMSO condition by ANC.

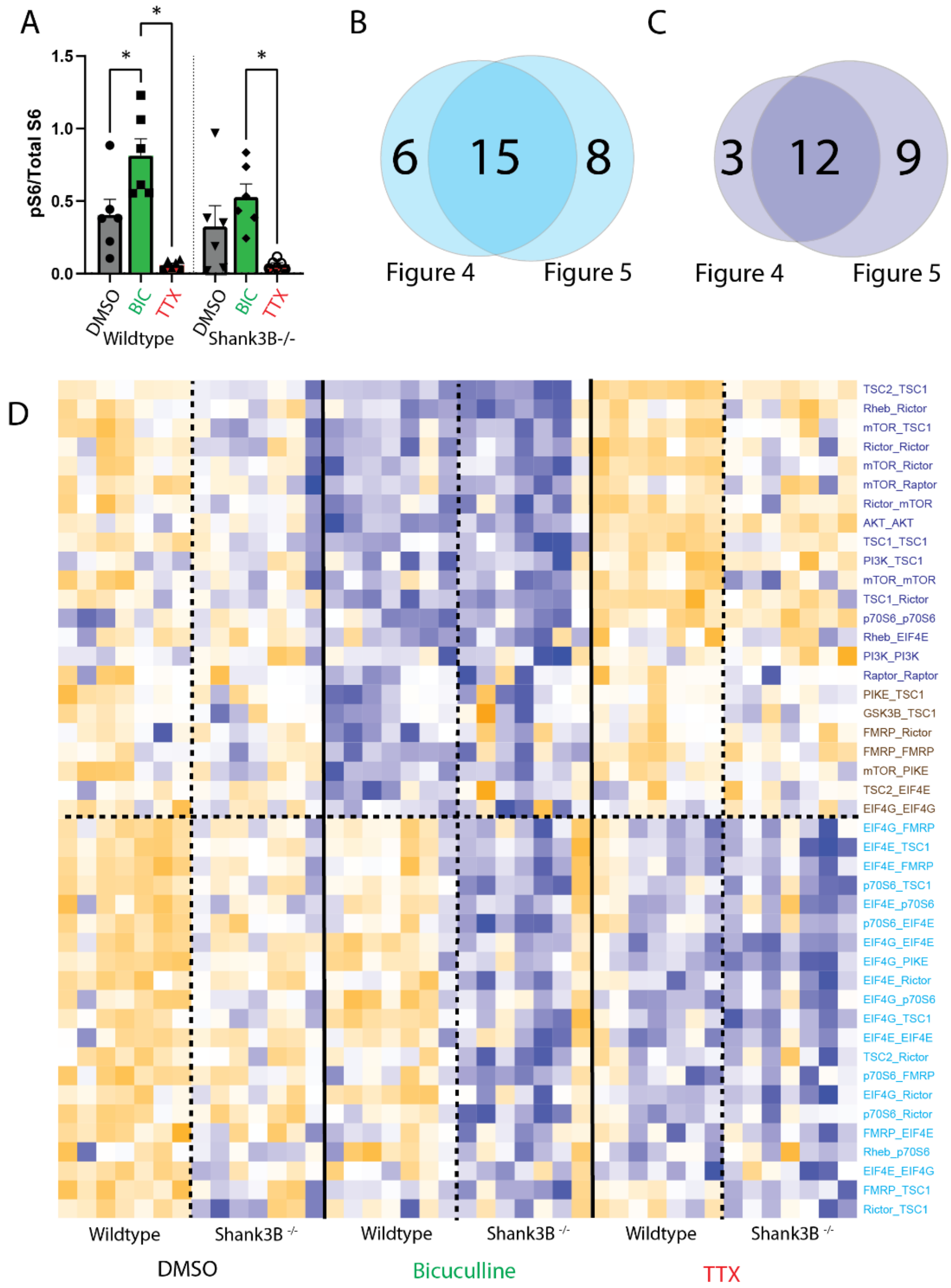


**Figure S3: Shank3 homeostatic scaling, Related to Figure5.** A) Quantification of western blots of phospho-to-total S6 shown in Fig 6A. * indicates $p<0.05$ compared to the DMSO condition (treatment effect) by 2-way ANOVA followed by Dunnett post-hoc testing. B,C) Venn diagram illustrating overlap between the BIC-only modules (B) and the “BOTH” modules (C) identified in the wild-type only scaling experiment (Figure 4) and the WT and Shank3 scaling experiment (Figure 5). D) Same heatmap as shown in Fig 5
